# Investigating Open Reading Frames in Known and Novel Transcripts using ORFanage

**DOI:** 10.1101/2023.03.23.533704

**Authors:** Ales Varabyou, Beril Erdogdu, Steven L. Salzberg, Mihaela Pertea

## Abstract

ORFanage is a system designed to assign open reading frames (ORFs) to both known and novel gene transcripts while maximizing similarity to annotated proteins. The primary intended use of ORFanage is the identification of ORFs in the assembled results of RNA sequencing (RNA-seq) experiments, a capability that most transcriptome assembly methods do not have. Our experiments demonstrate how ORFanage can be used to find novel protein variants in RNA-seq datasets, and to improve the annotations of ORFs in tens of thousands of transcript models in the RefSeq and GENCODE human annotation databases. Through its implementation of a highly accurate and efficient pseudo-alignment algorithm, ORFanage is substantially faster than other ORF annotation methods, enabling its application to very large datasets. When used to analyze transcriptome assemblies, ORFanage can aid in the separation of signal from transcriptional noise and the identification of likely functional transcript variants, ultimately advancing our understanding of biology and medicine.

## Introduction

Approximately 20,000 protein-coding genes have been annotated for the human genome ^1-5^. While a single isoform is often the source of the dominant protein ^6-8^, many human gene loci express isoforms that encode different protein sequences, some of which may be tissue-specific ^9, 10^. The NCBI RefSeq database, for example, contains an average of 6.9 isoforms for each human protein coding gene, which encode an average of 4.4 distinct protein sequences. The RefSeq annotations of the model organisms *C. elegans* and *A. thaliana* have on average 1.8 and 1.4 alternative isoforms with 1.3 unique protein variants respectively.

RNA sequencing technology has allowed an unprecedented look at the transcriptome in a wide variety of species, with multiple studies reporting large numbers of previously unknown transcripts for protein coding genes ^3, 11-14^. Consistent with previous reports about alternative splicing events ^15^, most of the novel transcripts reported in RNA-seq studies are observed in protein-coding regions ^16, 17^. Alternative splicing events can alter the translated protein through exon skipping, frame-shifting, and other changes ^19^. These events and their effects on translated proteins are an essential component of genome biology ^9^.

Changes in protein sequences may also be characteristic of disease states ^10, 20-23^ or of specific tissues ^9, 24, 25^. For example, splicing-induced changes in protein sequences have been associated with cancer development and progression, from activation of proto-oncogenes ^26^ to genome-wide splicing alteration in certain cancer types ^27, 28^. One example of why it is important to annotate all protein isoforms in the human genome is the widespread usage of exome sequencing in clinical settings. Exome capture methods have been extensively used to interrogate genetic variants and their associations with diseases, such as finding the genetic cause of a rare form of pediatric epilepsy ^29^, or identifying driver mutations in cancer ^30^. The technology is heavily dependent on the correct annotation of coding regions, and any exons that are un-annotated will simply be missed by exome studies.

However, many observed novel transcripts are likely to represent transcriptional noise ^18^; e.g., the original CHESS database assembled ∼29 million transcript variants from 10,000 RNA sequencing experiments, of which fewer than 2% were kept in the final annotation ^3, 4^. The ability to accurately identify non-functional isoforms can be a valuable tool in differentiating signal from noise in RNA-seq data, which is currently complicated by artifacts from computational methods as well as the amount of noise inherently present in the data ^18^.

Although many methods have been implemented for searching and assembling transcripts from RNA-seq data ^31, 32^, none of them identifies ORFs based on similarity to the original protein at the locus. Instead, previous approaches focused on identification of the longest ORF, typically requiring it to have the same start or stop codon positions ^33-35^, or on training machine learning models to find ORFs ^32, 36, 37^. None of these approaches consider the similarity of the resulting protein to other translations of the transcript.

In this study, we present ORFanage, a highly efficient and sensitive method to search for open reading frames in protein-coding transcripts, guided by reference annotation to maximize protein similarity within genes.

## Results

ORFanage utilizes protein-coding gene annotation by identifying ORFs in query transcripts that have the maximal sequence identity with a user-provided set of reference ORFs. This approach presumes that proteins produced by different transcripts at the same locus should be as similar as possible ^8, 38^. In our first set of experiments, we tested ORFanage’s ability to reconstruct the GENCODE and RefSeq protein-coding annotation given an annotation that includes one canonical ORF at each protein-coding gene locus. For these experiments, we used the MANE database to define the canonical ORFs, because MANE was created to be a “universal standard” of human protein-coding genes, and because both GENCODE and RefSeq contain every gene in MANE ^6^. These experiments illustrate how ORFanage can be used to make annotation more internally consistent.

As shown in Figure 1a and further discussed in the data filtering methods section below, many gene transcripts in both RefSeq and GENCODE are annotated with ORFs that differ from the canonical variant; e.g., 65% of ORFs in the RefSeq human annotation and 36% in GENCODE differ from the MANE ORF (Figure 1a). In principle, the presence of an ORF that differs from MANE does not imply an error; however, if another ORF can be found in the same transcript that has closer identity to MANE, then an error seems more likely. Furthermore, 8% of RefSeq and 43% of GENCODE transcripts in protein-coding loci have no ORFs annotated at all. By re-annotating each of the reference datasets using ORFanage, we identified numerous cases where a different ORF was more similar to the canonical protein. One example, from the *ZNF180* gene, is shown in Figure 1f.

**Figure 1.**
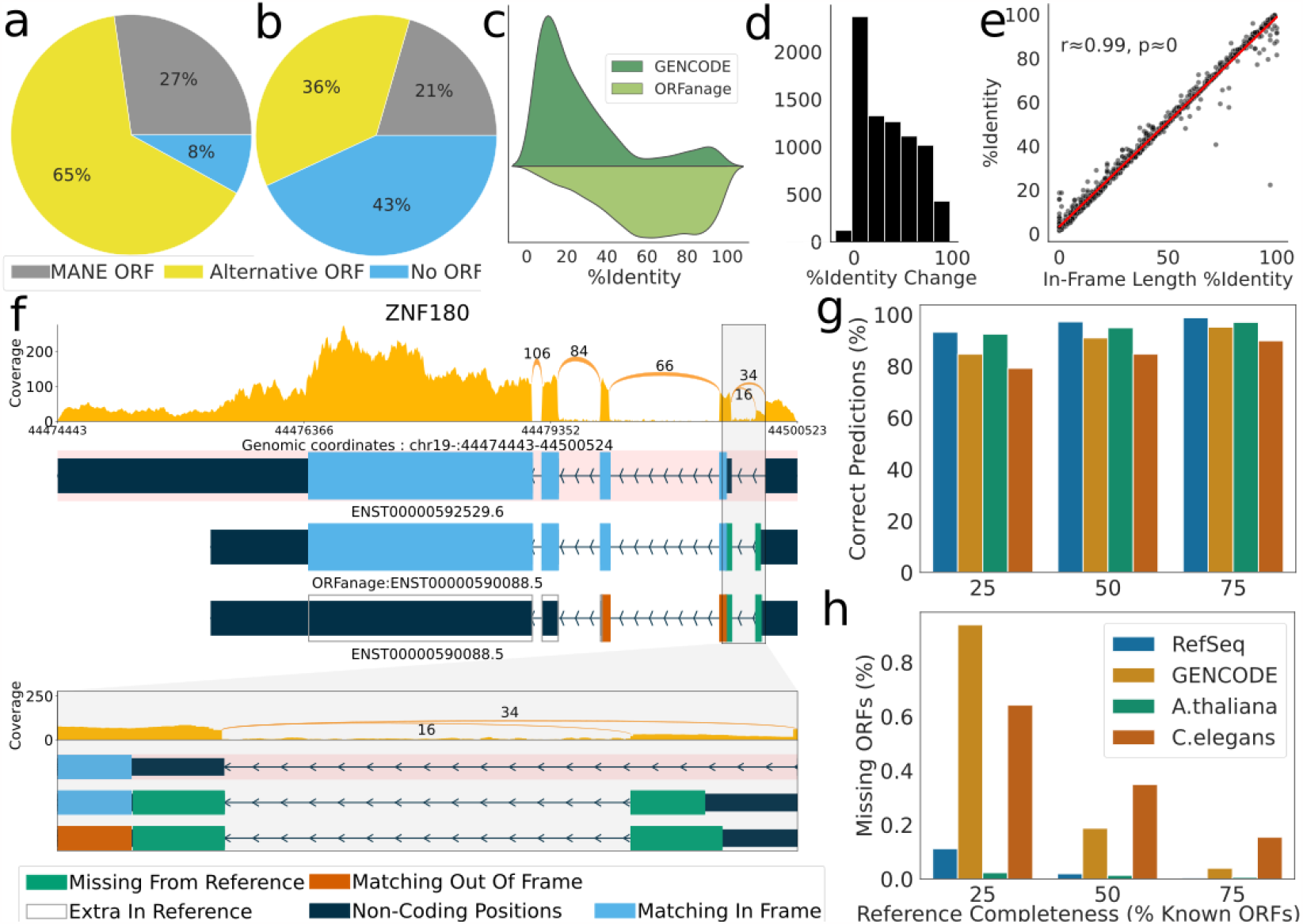
Overview of irregularities in reference database ORF annotation. a-b) Differences in ORFs at MANE loci as currently annotated for a) RefSeq and b) GENCODE annotations. Circular charts show, for each dataset, the proportions of transcripts annotated with the same ORF as MANE (grey), those with an alternative ORF not matching MANE (yellow), and transcripts in MANE loci that lack an annotated ORF (blue). c) Percent identity computed between the MANE protein and alternative ORFs as predicted by GENCODE (dark green) and ORFanage (light green). d) Histogram of the change in percent identity when replacing the GENCODE ORF with the ORFanage ORF. e) Correlation between percent identity computed via traditional alignment and In-Frame Length Percent Identity computed by ORFanage, illustrating the close similarity between the two metrics. f) A detailed look at alternative ORFs annotated by GENCODE and ORFanage for the ZNF180 gene. At top is the MANE isoform, shaded in pink, with its ORF shown in blue. Below it are two versions of an alternative isoform, with the ORFs annotated by ORFanage (middle) and GENCODE (bottom). Blue regions show where the protein sequence matches the MANE isoform, while green and orange show regions that are additional (green) or out of frame (orange) compared to MANE. At bottom is a zoomed-in view of the first intron and flanking ORF regions. g-h) Overview of the impact that completeness of reference annotation has on the accuracy of ORFanage. g) The percent of correctly inferred ORFs given different fractions of known reference ORFs for 4 organisms. h) Percentage of known ORFs that ORFanage failed to identify for different levels of reference completeness.

While we found that ORFs in a large majority of transcripts in the RefSeq human annotation were in agreement with those predicted by ORFanage (117,202 of 135,694) there were some striking differences, as illustrated in Table 1 and Supplemental Table 2. For example, we identified 2,122 transcripts in which an ORF annotated by RefSeq could be replaced by the canonical version from MANE without alterations. Similarly, 786 of the ORFs in the GENCODE human annotation could be replaced by their canonical variants from MANE. Because GENCODE and RefSeq both recognize MANE as a standard ^6^, it seems appropriate to choose the MANE ORFs over the alternative variants in these cases.

**Table 1.** Summary of differences between ORFs found by ORFanage and the originally annotated ORFs for all transcripts in RefSeq and GENCODE protein-coding genes. Comparisons to the MANE annotation refer to the ORFs from the MANE gene set, which is fully contained within both RefSeq and GENCODE.

|  |  |  |
| --- | --- | --- |
|  | Reference annotation |  |
|  | RefSeq | GENCODE |
| ORFanage finds the same ORF as reference | 117,202 | 63,966 |
| ORFanage finds a different ORF that matches MANE perfectly | 2,212 | 786 |
| ORFanage finds a different ORF, but reference matches MANE | 10 | 0 |
| No ORF annotated on reference transcript, ORFanage finds an ORF that matches MANE | 1,194 | 147 |
| No ORF annotated on reference transcript, ORFanage finds an ORF that is different from MANE | 9,240 | 35,393 |
| Other combinations | 5,836 | 27,994 |
| Total number of protein-coding transcripts | 135,694 | 128,286 |

We also found thousands of transcripts for which no ORF was listed, even though they were annotated under protein-coding genes and even though a valid ORF was identified by ORFanage. In GENCODE, we found an ORF that at least partially overlapped the MANE ORF in 35,540 out of 55,328 of these transcripts, including 147 transcripts that contained a perfect match to the MANE ORF. Although the RefSeq database had fewer protein-coding transcripts with no ORF listed, we still found 10,434 transcripts for which our method predicted a valid ORF, including 1,194 with a perfect match to MANE (Table 1, Supplemental Table 2).

We also looked at transcripts where both ORFanage and the reference annotation differed from MANE (5,301 in RefSeq and 7,957 in GENCODE). For these transcripts we computed the percentage of in-frame positions shared between the annotated proteins and the MANE protein, and observed that in 613 RefSeq and 7,005 GENCODE transcripts, ORFanage produced a protein that was closer to MANE (Figure 1c,d). In many cases the differences were minor, affecting only start coordinates or conserving different segments of the reference protein. In some cases, though, such as ZNF180 as shown in Figure 1f, ORFanage identified an ORF that conserved nearly all of the MANE protein sequence, while the protein encoded by the GENCODE ORF had no overlap with MANE.

When ORFanage found an ORF that differed from the one chosen by RefSeq or GENCODE, the ORFanage sequence had an equal or higher proportion of codons that matched MANE (Figure 1c-d), a property that is guaranteed by the algorithm. We confirmed these results by performing global alignments of the proteins to the MANE variants using EMBOSS Stretcher ^39^. The higher percent identity is a consequence of the metric that ORFanage maximizes, which we term In-frame Length Percent Identity (ILPI). To compute ILPI, our method first computes the total number of positions in an ORF that are in the same frame as the reference, thus coding for the same codons, which determines the In-frame Length (IL). Then ILPI is computed as the fraction of IL of the total length of the reference coding sequence. As illustrated in Figure 1e, the correlation between ILPI and percent identity computed via the Smith-Waterman algorithm is very high.

We then took a closer look at the 44,532 GENCODE transcripts where ORFanage found a different ORF. We found that ORFs identified by ORFanage often contained many novel positions (i.e., not matching MANE). More specifically, nearly 22% of positions in these novel ORFs are marked as potentially coding only by our method, and while many of these positions could be artifacts of partial transcript models included in the GENCODE annotation, some are likely to represent new functional variants of known proteins ^12, 22^.

The large number of missing annotations and overall observed improvements demonstrate the potential use of ORFanage at finding consistent ORFs in novel transcripts at protein-coding loci.

## Comparison to TransDecoder

TransDecoder, part of the Trinity package ^32^, can also find open reading frames in a set of transcripts, although it does not use reference annotation. Since its original release, the software has been adapted for use with transcript models that are assembled by programs such as StringTie ^40^ or Cufflinks ^41^.

We compared TransDecoder to ORFanage by using them both to re-annotate the protein-coding transcripts in the GENCODE human annotation (v41). We used default parameters for both methods, using the MANE annotation to guide ORFanage. Because TransDecoder produces multiple ORFs for some transcripts, we chose the longest to represent its output in those cases. We also removed any partial ORFs reported by TransDecoder.

Worth noting here is the considerably longer runtime required by TransDecoder, which took ∼41 minutes to annotate the GENCODE dataset while ORFanage required ∼20 seconds. If we had enabled the optional homology search option in TransDecoder, which would add further alignment steps, it would have taken far longer. The much greater computational requirement for TransDecoder could present a barrier in analyzing large collections of RNA-seq datasets.

For the 72,909 transcripts in GENCODE that contain a protein-coding sequence and that correspond to MANE loci, we compared the ORF predicted by ORFanage and TransDecoder to the ORFs annotated in GENCODE. ORFanage reproduced the GENCODE ORF for 88% (63,936) of the transcripts, while TransDecoder’s ORF matched GENCODE for 64% of the transcripts. TransDecoder entirely failed to find an ORF in 16,484 transcripts, compared to just 236 in which ORFanage failed to find an ORF.

There were 286 transcripts for which all three methods produced different ORFs. To evaluate these cases, we computed the percent identity between the predicted ORF and the corresponding MANE isoform. In 55% of the cases ORFanage predicted a protein more similar to MANE than the one predicted by TransDecoder, and in 45% of the cases the two methods produced variants with the same similarity to MANE.

Worth noting here is that TransDecoder is designed for finding ORFs *de novo*, without a need for guide annotation, and as such serves a different niche of applications than ORFanage; e.g., annotation of species for which no previous annotation is available.

## Impact of Reference Transcripts on Accuracy

In the next set of experiments, we set out to investigate how well our method can reconstruct a full set of protein sequences from subsets of reference data. We wanted to establish 1) how accuracy improves with an increase in the number of annotated ORFs at a locus and 2) the effects of choosing different subsets of known ORF variants on the accuracy of prediction. To answer the first question, we incrementally increased the number of ORFs provided to ORFanage as a reference. To address the second question, we repeated the experiment but randomly chose different sets of reference ORFs.

We repeated the iterative selection of reference transcripts 10 times, providing 25%, 50% and 75% of the reference ORFs as a guide each time. We ran our analysis on the human genome annotation as well as *A. thaliana* and *C. elegans* using the same protocol. For the human genome, we tested both RefSeq and GENCODE, because the two databases differ substantially in their ORF annotations. For each test run, we ensured that at least one transcript remained unannotated at each locus and that any non-coding transcripts were removed prior to the evaluation.

The diversity of transcripts annotated for *A. thaliana* and *C. elegans* is much lower compared to human reference annotations, with 1.8 and 1.4 transcripts per coding gene respectively, compared to 6 and 8 for RefSeq and GENCODE human annotations. Worth noting is that for *C. elegans*, only 4,422 suitable loci were identified based on the aforementioned criteria.

As expected, we observed an increase in accuracy as more reference annotation was provided. For the human genome, if we provided just a single reference ORF per locus (equivalent to 11% of all ORFs in RefSeq and 18% of all ORFs in GENCODE), ORFanage was able to perfectly re-create 85% of the RefSeq ORFs and 81% of GENCODE ORFs. When we provided 75% of the reference ORFs, ORFanage correctly re-created close to 99% of RefSeq and 95% of GENCODE ORFs (Figure 1g-h).

Even when ORFs were not identical to the original sources, the predictions produced by ORFanage were highly similar, averaging 81% for the non-identical predictions in the RefSeq dataset and 77% in GENCODE respectively.

Because *A. thaliana* and *C. elegans* have fewer annotated reference ORFs per locus, our random permutations had smaller effects on the results. Nevertheless, in *A. thaliana* ORFanage was able to correctly identify 91–97% of reference ORFs. For *C. elegans* the values were lower, ranging from 77% when a single random reference ORF was provided to 90% when guided by more complete annotations.

## Finding novel ORFs in assembled RNA-seq data

One of the main applications of ORFanage is to search for ORFs in datasets containing large numbers of transcripts that have not been assigned open reading frames. In particular, ORFanage can annotate transcriptome assemblies from RNA-seq experiments, which often contain many novel splice variants, even for well-annotated genomes ^3, 4, 42^. In these cases, ORFanage can identify valid ORFs for protein-coding transcripts using known annotation as a guide.

We next applied ORFanage to search for novel ORFs in experimental data, using data from the GTEx project ^43^, a high-quality collection of poly-A selected RNA-seq samples across multiple human tissue types. We focused our experiments on 1,448 samples from brain tissue because these represented the most diverse collection of samples in the dataset. We ran ORFanage on the complete, unfiltered set of assemblies containing 6,674,316 isoforms that were assembled originally for the CHESS human annotation database ^3, 4^.

We computed ORFs for all transcripts using the MANE annotation as the guide. For every MANE gene, we first identified all assembled transcripts overlapping that gene using gffcompare (similarity codes “=“, “c”, “k”, “m”, “n”, “j”, “e”), yielding 4,256,346 transcripts. We then computed the total gene expression for each transcript using the sum of TPM values for that transcript across all samples.

In our search for novel ORFs, we took a conservative approach: if a transcript could accommodate an ORF from either RefSeq or GENCODE, we assigned that ORF to the transcript. Additionally, we removed ORFanage predictions for all transcripts marked as non-coding by either RefSeq or GENCODE. Because multiple distinct transcripts can contain the same novel ORF, we simplified our analysis by computing the total TPM aggregated across transcripts sharing the same ORF. In transcripts for which no ORF was assigned, we computed the total TPM as the sum of TPMs for that transcript across all samples. This selection left us with a total of 3,046,286 novel transcripts representing 1,006,547 ORF variants.

Next to focus on highly expressed cases, we considered 4,190 loci where more than 50% of expression came from novel transcripts and ORFs (Figure 2a). Many of the transcripts at these loci either had no valid ORF or else contained an ORF that was highly dissimilar from the canonical MANE protein. We therefore narrowed our focus to 462 loci where over 50% of expression was due to a single novel ORF. Of those, only 24 loci (Supplementary Table 1) were at least 70% identical to the MANE protein and had cumulative expression greater than 1000 TPM across all samples (Figure 2b-d, Supplemental Figure 1-2). For instance, in the PLGLB gene, an exon skipping event via a novel intron leads to the loss of the original start codon and a different, slightly longer N-terminal amino acid sequence. Interestingly, we observed very similar exon skipping events in two different paralogs of this gene, PLGLB1 and PLGLB2, shown in Figure 2c-d. In both cases, the alternative protein contains a different initial coding exon that replaces exon 1 of the MANE isoform, and in both cases, the majority of the expression comes from the alternative (non-MANE) isoform, suggesting that the MANE isoforms are not the dominant ones.

**Figure 2.**
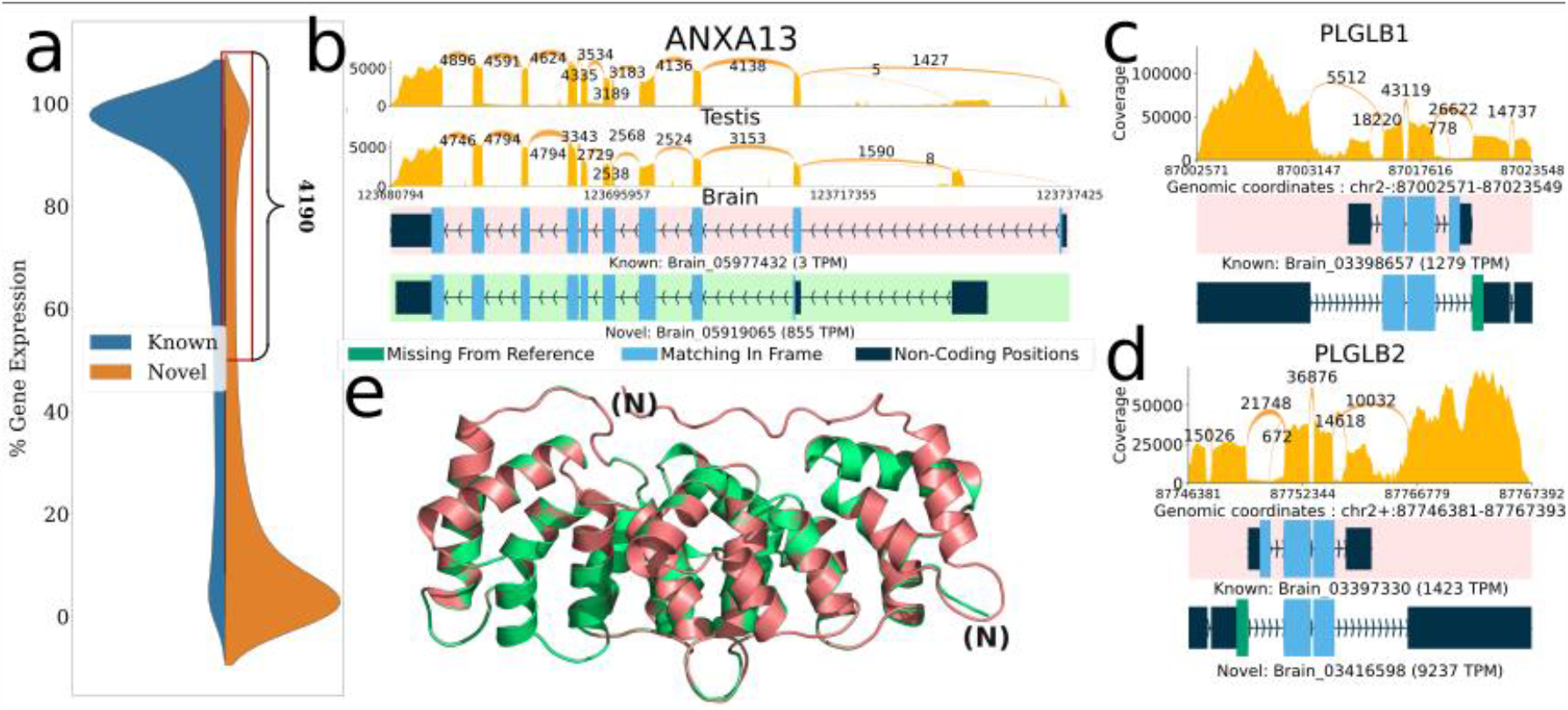
Novel ORFs in the GTEx dataset inferred using ORFanage. a) Overall distribution of loci by percent gene expression (y-axis) that comes from novel (orange) and known (blue) transcripts and a zoomed-in view of the region containing 4,190 loci where >=50% of the total expression comes from transcripts with novel ORFs or novel transcripts without an ORF. b,c,d) Sashimi plots illustrating selected examples of novel ORFs that were identified by ORFanage, each depicting a different type of variation. In each plot, coverage and splice junction values are cumulative across all samples ^44^. The uppermost transcript, highlighted with a pink background, shows the MANE annotation. Expression levels measured in TPM are shown for each transcript. b) An alternative 5’ exon in ANXA13 that changes the start codon and shortens the ORF. e) The 3D alignment of the MANE protein (pink) to the novel ORF (green) computed by Alphafold2 and visualized via PyMOL ^45^ is shown below with the N-termini labeled for each. c,d) Two plots show similar novel ORFs for two paralogous genes PLGLB1 and PLGB2, where skipping of the 1st reference coding exon is effectively offset by the introduction of an upstream novel exon with an alternative start codon.

Another striking example of a novel ORF among these 24 loci occurs in the ANXA13 gene (Figure 2b,e), which is a member of the family of annexin genes responsible for the production of calcium-dependent membrane-binding protein variants ^46^. Proteins in this family contain two major domains, one at the C-terminus for the Ca^2+^ binding effect, and the other at the N-terminus responsible for the membrane interactions. While the core domain at the C-terminus is highly conserved across the gene family, the N-terminus is variable ^47^, allowing for tissue-specific regulation ^48, 49^ and localization ^50^.The two known forms of the gene differ only in the length of the last helical structure, where the incorporation of additional peptides allows for an extension of the first helix.

In our results, the majority of expression of ANXA13 came from a novel variant of the gene characterized by a mutually exclusively alternative splicing event which results in the switching of the start-codon-harboring exon for another one downstream, as shown in Figure 2b. The novel variant has an alternative methionine, followed by a glycine, that serves as its start codon, preserving much of the protein sequence with a new N-terminus. We also observed that this isoform was dominant in brain tissue, while the MANE isoform was dominant in testis (and other tissues).

We investigate how the change would impact the protein’s structure by folding it with AlphaFold2 via ColabFold ^51, 52^. We observed an increase in the pLDDT score from 94 to 97, suggesting an even more stable structure for the new isoform, due to the removal of an unstructured segment at the N-terminus of the MANE isoform (Figure 2e). The alternative protein identified here matches a variant that was previously annotated as the third isoform of AXNA13 in *Pan troglodytes* ^53^ and *Papio anubis* ^54^.

## Conclusion

Our understanding of the transcriptional complexity of eukaryotic genomes has expanded dramatically over the years, yet the full extent and functional implications of alternative splicing are not yet entirely understood. A comprehensive evaluation of the proteome generated by alternative splicing is critical not only for identifying anomalies in disease states but also for identifying novel protein variants with distinct functions.

Our experiments demonstrate the effectiveness of ORFanage for identifying open reading frames in a set of transcripts by using reference annotation as a guide. ORFanage can recover most of the original annotation of the human genome using any of several widely used annotation databases, and it can also identify inconsistencies in those databases. More specifically, we showed that it can identify cases where an annotated ORF can be adjusted so that it matches a canonical protein sequence more closely than the original annotation from either GENCODE or RefSeq. While not explicitly tested here, our protocol can be easily combined with programs like Liftoff ^55^ to facilitate comprehensive annotations of genomes of various ancestries that include not only transcripts but coding regions as well.

ORFanage can be used in conjunction with RNA-seq alignment and assembly to identify ORFs in novel transcripts, and to guarantee that those ORFs match the reference annotation as closely as possible. Whether using long-read alignments directly or assembled transcripts, this approach can uncover valuable insights into isoforms within protein-coding regions, leading to a better understanding of their effects on biological systems. And because RNA-seq datasets often produce large numbers of novel transcripts, the efficiency and scalability of ORFanage make it suitable for datasets of any size. We have recently applied our method to annotate ORFs in novel transcripts for the revised CHESS 3 ^4^ catalog, and to help identify novel structurally stable isoforms that were then confirmed using AlphaFold2 ^56^.

ORFanage can also be a valuable aid to isolating true signal from noisy transcriptome data. Assuming that proteins produced from alternative transcripts need to remain similar in order for genes to function correctly ^56^, the ORF structures in the observed isoforms should be similar as well. Our approach can identify transcripts that cannot accommodate a similar ORF to the reference, serving as a noise filtering step in RNA-seq analysis.

## Methods

ORFanage is based on the direct comparison of intervals that make up the exonic structures of query and reference transcripts. This optimization technique does not require sequence alignment or pre-computed genome indices, greatly reducing the computational burden of running the tool and making the analysis far more efficient than an alignment-based approach. We have tested ORFanage on datasets comprising tens of millions of transcripts assembled from thousands of RNA-seq experiments ^3, 4^ and found that it runs robustly on these data.

## Creating Bundles of Transcripts

ORFanage operates on “bundles” of data, defined as the union of a set of overlapping reference ORFs with a set of query transcripts that overlap 1 or more of the reference ORFs. In order to reduce the impact of annotation errors such as readthrough transcription, ORFanage only loads CDS coordinates for each reference transcript, discarding non-coding exonic coordinates.

Once both the reference and query datasets are loaded into memory and sorted internally, bundling is done in linear time by iterating over transcripts and collecting groups of all overlapping transcripts. This technique is insensitive to any information on gene boundaries, and readthrough transcription, commonly present in RNA-seq assemblies, may lead to several genes being combined into a single locus. In some cases, genes may genuinely overlap, and in such instances ORFanage might compare the ORFs of unrelated genes, possibly leading to incorrect inferences. To combat this problem, ORFanage gives users the option to group transcripts by gene IDs.

## Interval Comparison

For each query transcript in a bundle, ORFanage performs a comparison to each reference CDS. For each pair being compared, an intersection is computed to identify all intervals that belong to both the query and the reference. The process is performed for all reference transcripts and duplicate intervals are removed.

After a set of candidate overlaps is identified, ORFanage continues to search for the optimal start and end coordinates for each interval, discarding any incomplete ORFs in the process. We define a valid open reading frame as an uninterrupted sequence of 3-base codons that begins with a start codon (usually ATG in humans), ends with a stop codon (TAA, TAG, and TGA in humans), and does not contain any other stop codons other than the final one. While only one valid stop codon can be found by extending any given ORF, multiple start (ATG) codons may be present in a single ORF. In ORFanage, an optimal start codon is the one that maximizes the number of bases which are in the same frame as the reference ORF while minimizing the number of coordinates that do not match or that match out-of-frame (Figure 3).

**Figure 3.**
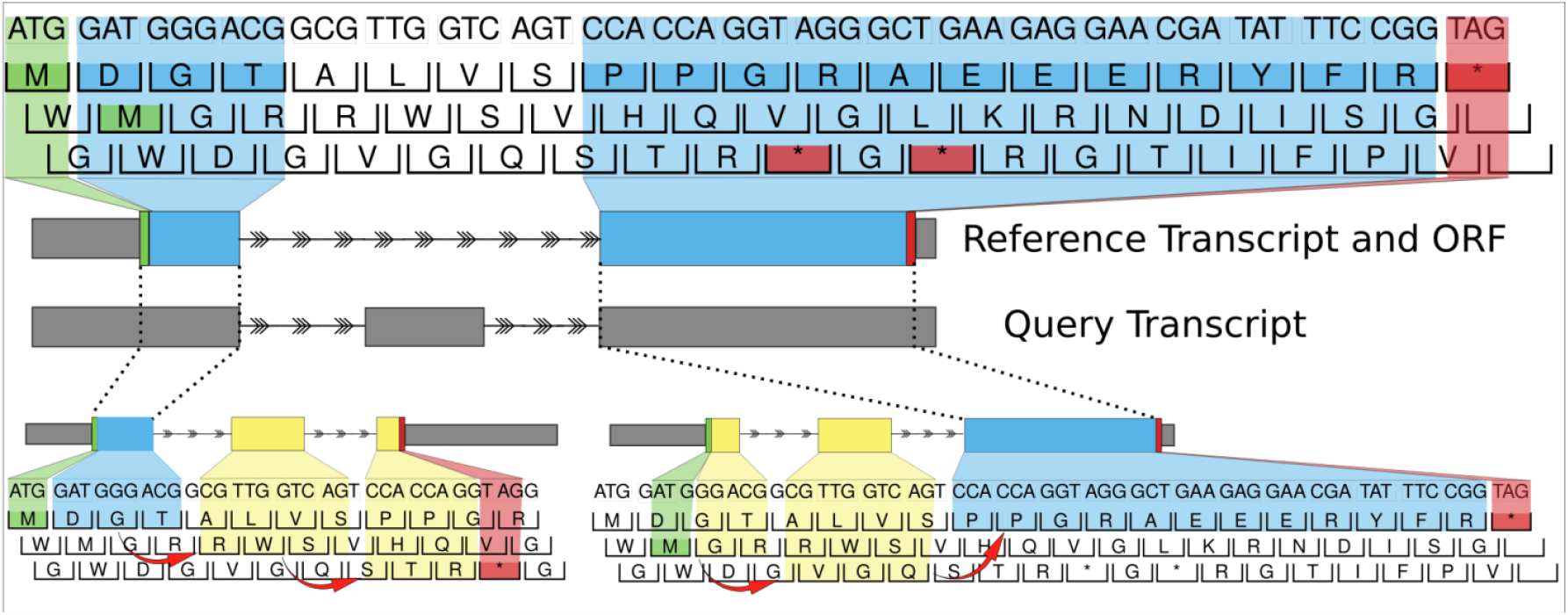
Diagram illustrating the algorithm implemented in ORFanage. ORFanage begins by computing overlaps between reference ORF and query transcript. In the figure, dashed lines are used to connect matching intervals. For each overlap it extends coordinates towards the 3’ and 5’ ends based on suitable parameters. During extension, any changes to the exon structure may introduce shifting of the original frame (as indicated by red arrows). Once all intervals have been evaluated, ORFanage compares the results and reports the one with the highest score. In the figure, matching residues to the reference are highlighted in blue, and mismatching residues are highlighted in **yellow**. In this example, ORFanage selects the longer ORF on the lower right, which has 13 out of 17 matching residues, compared to the ORF on the lower left with only 4 out of 17 matching residues.

After all intervals have been examined, if multiple distinct ORFs are plausible, ORFanage performs a heuristic selection of the optimal ORF based on a series of configurable steps. Internally, for every unique ORF, the software computes three scores which are applied successively to each set of candidate ORFs to find the best result:

1. the Inframe Length (IL), defined as the number of positions that are shared with the reference in the same coding frame,

2. ILPI, defined as the fraction of IL with respect to the length of the reference ORF, and

3. the length of the ORF.

When maximizing ILPI, ORFanage will prioritize ORFs that have as little novel sequence present as possible, where “novel” is defined as sequence that is not present in the reference ORF. When maximizing IL instead, ORFanage might select longer ORFs with more novel sequence if that choice increases the number of matches with the reference.

## Additional parameters

As shown in our analysis, the ILPI metric is an effective function to assess which ORF to pick for a given isoform and corresponds closely to percent identity. It is not identical to the familiar percent identity measure, which would be more expensive to compute. For applications that might require it, ORFanage includes support for computing a Smith-Waterman alignment between the reference ORF and the ORFs identified by ORFanage, as part of the final validation of the open reading frames. ORFanage also includes an option to measure evolutionary conservation of any ORF by computing PhyloCSF scores. This option is implemented via an integrated PhyloCSF++ module ^57, 58^. Finally, ORFanage contains a multi-threading option, under which it can process each bundle in parallel, further speeding its runtime. In our tests, ORFanage was able to process 4,256,346 collected from 1,448 brain samples of the GTEx dataset, using the MANE annotation as a reference, in 7 minutes using 24 cores of an Intel Xeon 6248R 3GHz processor, with all other parameters set to defaults. A random individual sample from the same dataset (SRR598396) was processed in 8 seconds.

## Datasets

Studies of the human genome account for a large proportion of transcriptomic data being generated today, and several annotation databases are available for these studies. For our evaluation of ORFanage on the human genome, we used both the RefSeq (release 110) and GENCODE (release 41) annotations ^1, 2^.

To investigate the utility of ORFanage on other organisms, we focused on the well-studied *A. thaliana* and *C. elegans* genomes, both of which have highly curated annotations of the transcriptome and proteome. Since for each of these two genomes only a single reference annotation was available, we chose to investigate how well ORFanage could reconstruct the ORFs using a bootstrapping technique, which allowed us to evaluate the concordance of annotated ORFs with the ones inferred by ORFanage.

For our evaluations on GTEx data, we used 1,448 poly-A selected RNA-seq samples representing 13 brain regions (age ≥ 20) from GTEx release 7 ^43^. Samples were aligned with HISAT2 ^59^, assembled with StringTie 2 ^31^, and merged with gffcompare. Coverage and splice junction summaries were extracted using the TieBrush suite ^44^.

## Data preparation

While ORFanage can handle several types of exceptions to the normal rules governing ORFs, such as alternative (non-ATG) start codons, selenoproteins, and otherwise overlapping genes, for our evaluations we removed these exceptions in order to measure accuracy on genes that conform to standard rules.

We began by choosing a set of genes to be used as a reference for human annotation. The MANE database ^6^ was created by the developers of RefSeq and GENCODE as a resource of human genes where both databases agree precisely on the complete exon-intron structure as well as on the coding sequence of every gene in the database. MANE contains one canonical transcript for nearly every protein coding gene, plus a small number (62 in release 1.0) of medically-relevant transcripts that differ from the canonical ones. In our reference set, we included all genes in MANE except for 1) genes with non-ATG start codons, 2) selenoproteins, and 3) polycistronic genes. For our evaluations of both RefSeq and GENCODE, we retained only transcripts corresponding to the remaining MANE genes.

In some cases, manual curation might have altered RefSeq or GENCODE to create unusual ORFs. For instance, some partial transcripts have been manually curated to show usage of an alternative start codon, despite other ORFs at the locus containing a canonical start codon. Because we do not know whether such exceptions are intentional, we decided to avoid penalizing RefSeq or GENCODE and filter out such cases, as follows. First, we used gffread ^60^ to identify and remove all transcripts that did not contain valid start and stop codons. Second, we searched for all pairs of overlapping ORFs that were labeled with different gene IDs and removed all such occurrences. In addition, for the RefSeq dataset we also removed 846 genes that had transcripts with known exceptions as annotated by NCBI. In the end, our filtering resulted in the removal of 1,423 genes out of 20,442 genes from RefSeq (release 110) and 1,869 genes out of 20,427 from GENCODE (version 41).

For the *A. thaliana* and *C. elegans* annotation datasets ^61, 62^, we used the primary model organism annotation as the reference, after filtering out genes with non-ATG start codons, selenoproteins, and polycistronic genes.

## Code availability

The core method is implemented in C++ and based on the GFFutils ^60^ and KSW2 ^63, 64^ libraries. The code and test data are available for download on GitHub: https://github.com/alevar/ORFanage. Version 1.0 of the software was used in this study. Jupyter notebooks used to generate all results described in the manuscript are provided separately at https://github.com/alevar/ORFanage_tests.

## Acknowledgements

We would also like to thank Christopher Pockrandt for helpful discussions on phyloCSF++ implementation and usage.

## Author Contributions

A.V. and B.E. conceived and developed the original idea. A.V. developed and implemented the final method and experiments. A.V., B.E., S.L.S and M.P. conceptualized the study, methods, and wrote the manuscript.

