## Supplemental Materials for "Investigating Open Reading Frames in Known and Novel Transcripts using ORFanage"

| **Supplemental Table 1.** List of 24 loci containing novel ORFs that have high expression levels. The data was collected from the subset of GTEx samples comprised of brain tissue. | | | | |
| --- | --- | --- | --- | --- |
| Gene Name | MANE TX | Novel TX | Gene Expression Novel % | Gene Expression (TPM) |
| GOLGA8B | ENST00000683415.1 | Brain_01934434 | 105755 | 53 |
| ANLN | ENST00000265748.7 | Brain_05557376 | 70084 | 76 |
| TUBGCP6 | ENST00000248846.10 | Brain_04137050 | 60672 | 60 |
| NKX6-2 | ENST00000368592.8 | Brain_00661433 | 51226 | 50 |
| AKAP3 | ENST00000228850.6 | Brain_01269902 | 46401 | 96 |
| SLC2A4RG | ENST00000266077.5 | Brain_03889109 | 32793 | 71 |
| ANK1 | ENST00000289734.13 | Brain_05914479 | 31276 | 50 |
| CD22 | ENST00000085219.10 | Brain_03028658 | 24579 | 68 |
| HOOK2 | ENST00000397668.8 | Brain_03025283 | 20813 | 55 |
| ABCA7 | ENST00000263094.11 | Brain_03022976 | 17695 | 57 |
| PLGLB2 | ENST00000359481.9 | Brain_03416598 | 12096 | 76 |
| PLGLB1 | ENST00000355705.4 | Brain_03431431 | 11908 | 82 |
| PARVG | ENST00000444313.8 | Brain_04138511 | 7773 | 56 |
| RFX7 | ENST00000559447.8 | Brain_01934679 | 7426 | 55 |
| SHROOM3 | ENST00000296043.7 | Brain_04680479 | 6575 | 54 |
| C19orf71 | ENST00000329493.6 | Brain_03023118 | 4488 | 61 |
| NEXMIF | ENST00000055682.12 | Brain_06469912 | 4433 | 86 |
| ZNF628 | ENST00000598519.2 | Brain_03024624 | 4068 | 52 |
| LEFTY1 | ENST00000272134.5 | Brain_00005009 | 3386 | 63 |
| GUCA1B | ENST00000230361.4 | Brain_05238473 | 3214 | 77 |
| CALML6 | ENST00000307786.8 | Brain_00000078 | 2317 | 72 |
| KLK5 | ENST00000336334.8 | Brain_03030776 | 2176 | 67 |
| SGO2 | ENST00000357799.9 | Brain_03394864 | 1999 | 57 |
| ANXA13 | ENST00000419625.6 | Brain_05919065 | 1502 | 56 |

| **Supplemental Table 2**. Details of the similarities and differences in ORFs found in protein-coding genes among RefSeq, GENCODE, and ORFanage. The upper rows compare ORFanage and RefSeq, and the lower rows compare ORFanage and GENCODE. We provide a comparison of ORFs that match MANE and do not match MANE in each comparison. For example, the number of RefSeq transcripts that have a non-MANE ORF but where corrected ORFanage found a perfect match to MANE ORF was 2212. Similarly, the number of non-MANE ORFs for which both RefSeq and ORFanage agree is 80,180. In the table, "=" means the ORFs agree perfectly, and "≠" means they do not match perfectly. "No ORF" counts transcripts for which no ORF was annotated. | | | | | | |
| --- | --- | --- | --- | --- | --- | --- |
|  |  | **ORFanage** | | | | |
|  |  | matches MANE | does not match MANE | | No ORF | Total |
| **RefSeq** | matches MANE | 37022 | 11 | | 0 | 37033 |
|  | does not match MANE | 2212 | **Total** | 85470 | 83 | 87765 |
|  |  |  | **RefSeq = ORFanage** | 80180 |  |  |
|  |  |  | **RefSeq ≠ ORFanage** | 5290 |  |  |
|  | No ORF | 1194 | 9240 | | 462 | 10896 |
|  | Total | 40428 | 94721 | | 545 | 135694 |
| **GENCODE** | matches MANE | 26283 | 23 | | 0 | 26306 |
|  | does not match MANE | 787 | **Total** | 45617 | 248 | 46652 |
|  |  |  | **GENCODE = ORFanage** | 37683 |  |  |
|  |  |  | **GENCODE ≠ ORFanage** | 7934 |  |  |
|  | No ORF | 147 | 35393 | | 19788 | 55328 |
|  | **Total** | 27217 | 81033 | | 20036 | 128286 |

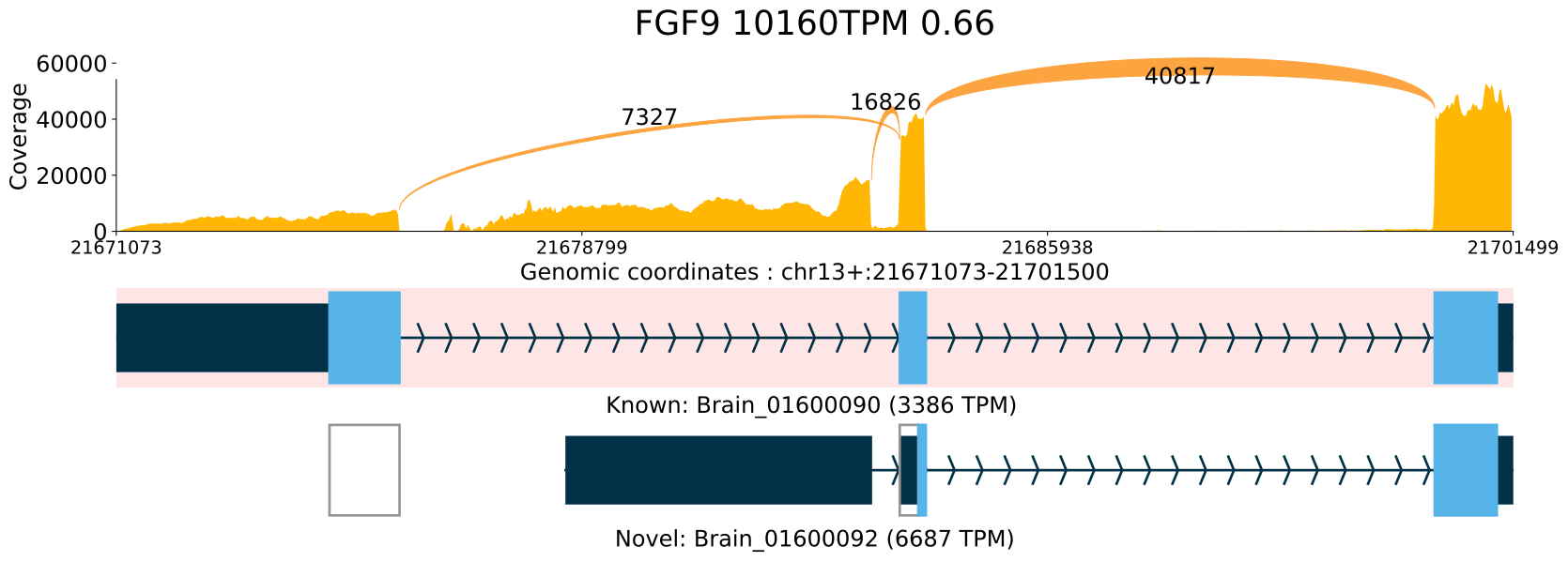
**Supplemental Figure 1.** Sashimi plot of the FGF9 locus illustrating a comparison of the canonical ORF to the novel ORF found by ORFanage in a highly expressed isoform. Coverage and splice junction values are cumulative across all samples. The uppermost transcript, highlighted with a pink background, always shows the MANE annotation. Expression levels measured in TPM are shown for each transcript. In FGF9 gene, we have observed the majority of expressed transcripts (66%) to be assembled with a novel first exon, resulting in the loss of the original start codon and likely introducing a new start of translation which preserves most of the original protein sequence.

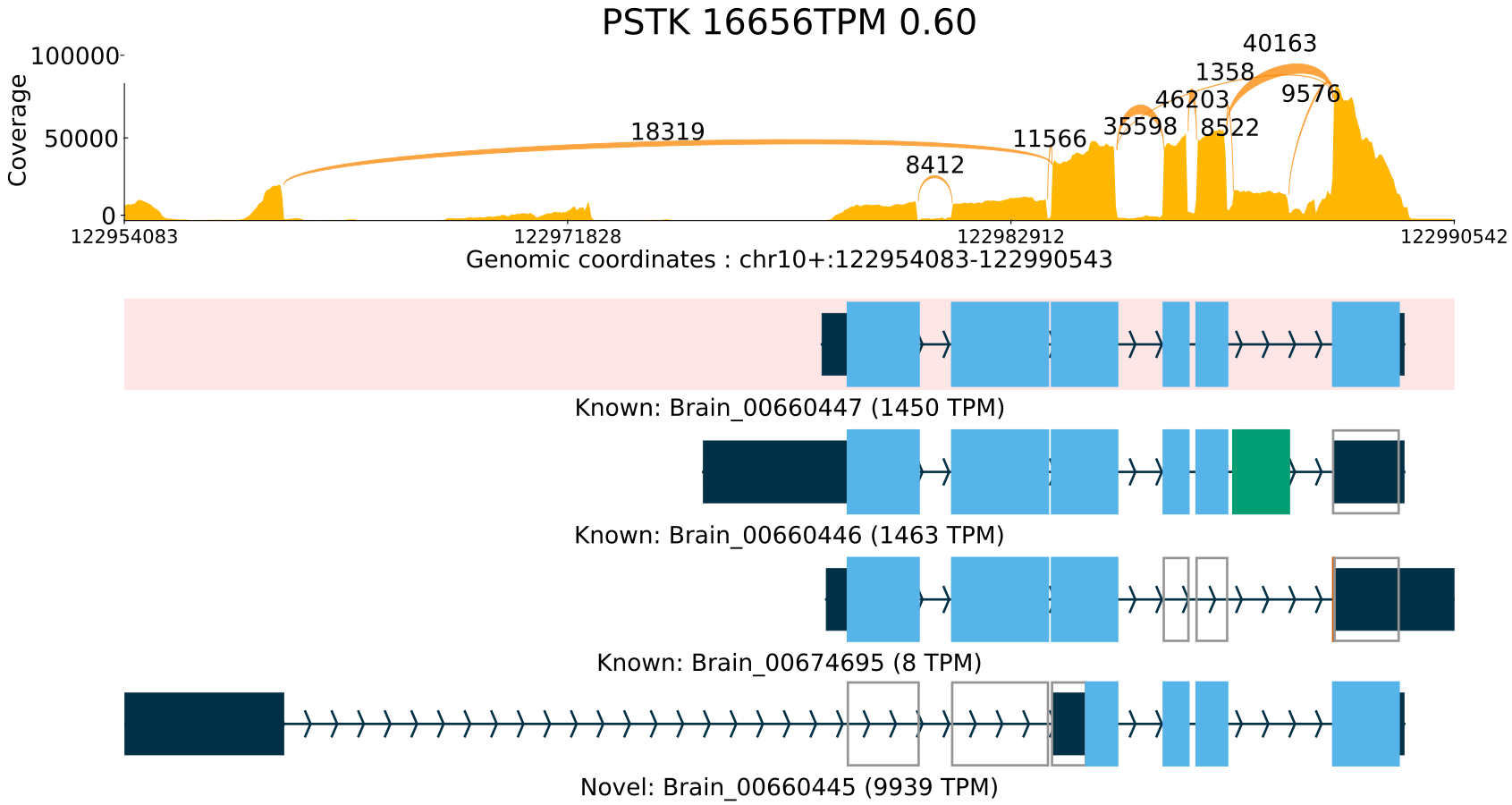
**Supplemental Figure 2.** Sashimi plot of the PSTK locus illustrating a comparison of the canonical ORF to the novel ORF found by ORFanage in a highly expressed isoform. Coverage and splice junction values are cumulative across all samples. The uppermost transcript, highlighted with a pink background, always shows the MANE annotation. Expression levels measured in TPM are shown for each transcript. In this locus an exon skipping event via a novel intron leads to the loss of the original start codon and truncation of the main protein.
